# The hippocampus as the switchboard between perception and memory

**DOI:** 10.1101/2020.05.20.104539

**Authors:** Matthias S. Treder, Ian Charest, Sebastian Michelmann, María Carmen Martín-Buro, Frédéric Roux, Fernando Carceller-Benito, Arturo Ugalde-Canitrot, David T. Rollings, Vijay Sawlani, Ramesh Chelvarajah, Maria Wimber, Simon Hanslmayr, Bernhard P. Staresina

## Abstract

Adaptive memory recall requires a rapid and flexible switch from external perceptual reminders to internal mnemonic representations. However, owing to the limited temporal or spatial resolution of brain imaging modalities used in isolation, the hippocampal-cortical dynamics supporting this process remain unknown. We thus employed an object-scene cued recall paradigm across two studies, including intracranial Electroencephalography (iEEG) and high-density scalp EEG. First, a sustained increase in hippocampal high gamma power (60-110 Hz) emerged 500 ms after cue onset and distinguished successful vs. unsuccessful recall. This increase in gamma power for successful recall was followed by a decrease in hippocampal alpha power (8-12 Hz). Intriguingly, the hippocampal gamma power increase marked the moment at which extrahippocampal activation patterns shifted from perceptual cue towards mnemonic target representations. In parallel, source-localised EEG alpha power revealed that the recall signal progresses from hippocampus to posterior parietal cortex and then to medial prefrontal cortex. Together, these results identify the hippocampus as the switchboard between perception and memory and elucidate the ensuing hippocampal-cortical dynamics supporting the recall process.

**Significance:** How do we adaptively switch from perceiving the external world to retrieving goal-relevant internal memories? To tackle this question, we used – in a cued-recall paradigm - direct intracranial recordings from the human hippocampus complemented by high-density scalp Electroencephalography (EEG). We found that a hippocampal signal ~500 ms after a perceptual cue marks the conversion from external (perceptual) to internal (mnemonic) representations. This sets in motion a recall cascade involving posterior parietal and medial prefrontal cortex, revealed via source-localised and time-resolved EEG alpha power. Together, these results unveil the hippocampal-cortical dynamics supporting rapid and flexible memory recall.

## Introduction

Imagine spotting a familiar face at a (real) conference. As your acquaintance approaches, you frantically try to recall the last time the two of you met and – without sneakily glancing at the nametag – remember what their name was. This example illustrates how adaptive behaviour often requires us to shift our focus from external sensory information to internal mnemonic representations. In experimental terms, this scenario constitutes a cued recall task, where a reminder cue may or may not trigger recall of associated mnemonic target information. How does our brain accomplish the feat of converting an external reminder into a target memory?

According to computational models, the hippocampus links disparate cortical representations into a coherent memory trace (Lisman, 1999; Wallenstein et al., 1998). It retains pointers to the cortical sites involved in the initial experience (Goode et al., 2020; Teyler and DiScenna, 1986) such that presenting a partial reminder prompts reinstatement of the entire association via hippocampal pattern completion (Marr, 1971; Norman and O’reilly, 2003). In support of these models, human functional magnetic resonance imaging (fMRI) studies linked hippocampal activation with cortical reinstatement of mnemonic target representations during successful recall (Bosch et al., 2014; Gordon et al., 2013; Grande et al., 2019; Horner et al., 2015; Ritchey et al., 2013; Staresina et al., 2013; Staresina et al., 2012b). However, the relatively poor temporal resolution of the fMRI signal leaves open whether the hippocampus precedes or follows mnemonic reinstatement, let alone whether hippocampal engagement would mark the rapid switch from perceptual cue to mnemonic target representations.

Moreover, the cognitive complexity and representational richness of memory recall likely requires concerted engagement of wider brain networks (Olsen and Robin, 2020; Ranganath and Ritchey, 2012). Indeed, beyond the hippocampus, neuroimaging work has consistently implicated a particular set of cortical regions in episodic memory tasks (Ritchey and Cooper, 2020; Rugg and Vilberg, 2013), herein referred to as the ‘cortical retrieval network’ (CRN). The CRN overlaps with the ‘default mode network’ (Buckner et al., 2008) and includes posterior parietal regions as well as medial prefrontal cortex. It has been linked to retrieval success across multiple stimulus domains (Hayama et al., 2012) as well as to episodic (re)construction processes (Benoit and Schacter, 2015; Hassabis and Maguire, 2007). Critically, a recent study employing ‘lesion network mapping’ suggests that the hippocampus serves as a functional hub linking these cortical nodes in service of memory processes (Ferguson et al., 2019). While these results indicate that successful memory relies on intricate hippocampal-cortical interactions, the temporal dynamics within the CRN are challenging to resolve with fMRI alone, hampering understanding of different CRN regions’ contributions (Ritchey and Cooper, 2020).

To overcome these limitations, we used intracranial Electroencephalography (EEG) complemented by high-density scalp EEG to reveal (i) the role of the hippocampus in the conversion of perceptual cues to mnemonic targets and (ii) the ensuing dynamics in the fronto-parietal retrieval network.

## Results

### Behaviour

We used the same memory paradigm (Figure 1) in an intracranial EEG (iEEG) study (n = 11) and a high-density scalp EEG study (n = 20). In addition, we conducted ‘localiser’ runs (Figure 1A) to train a classifier to distinguish brain patterns of object vs. scene representations (see below). In the memory experiment (Figure 1B), participants were presented with pairs of object and scene images during encoding. During retrieval, a cued recall task was employed in which only one of the images was shown (‘cue’), with the question whether the associated image (‘target’) was also remembered. Catch trials were interspersed in which participants were prompted to describe the target image after giving a “Remember” response.

**Figure 1.**
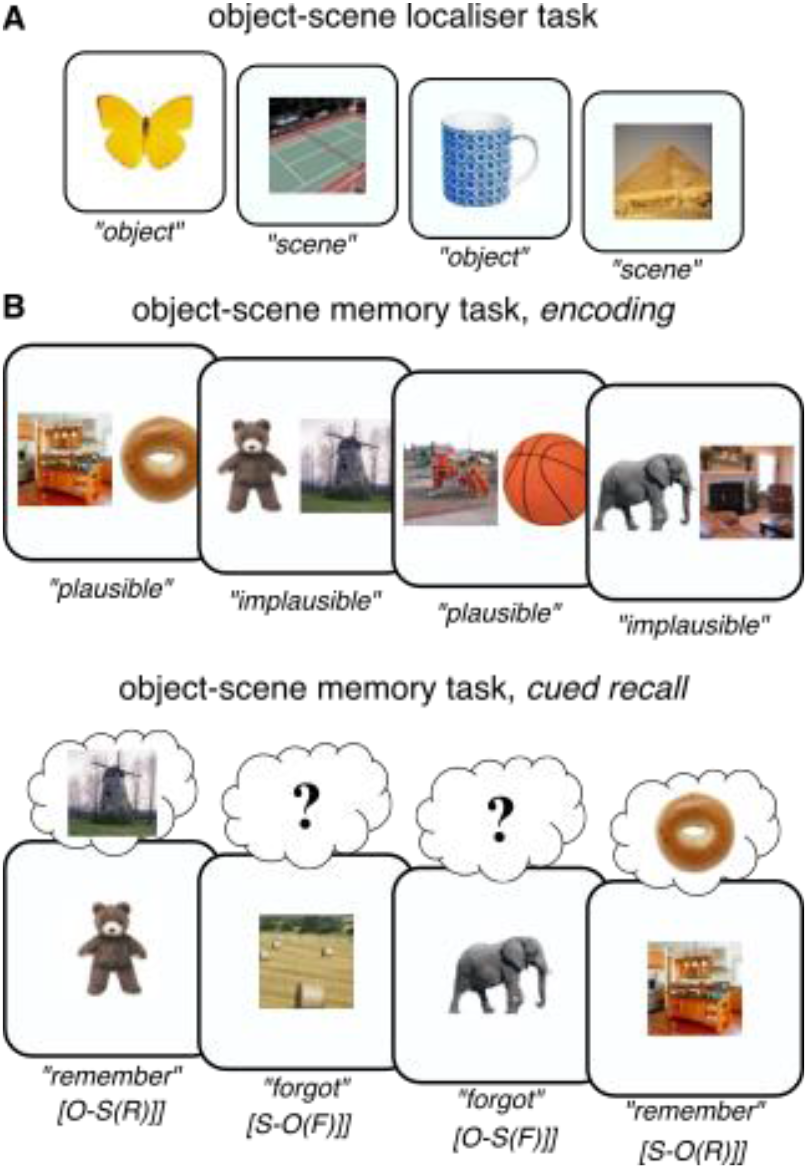
Experimental paradigm. **A**. In a *localiser* session, participants saw trial-unique images of objects and scenes and indicated the category of the given image. This part served as an independent training dataset for multivariate pattern analyses. **B**. The main experiment employed an object-scene memory task, consisting of an encoding phase (*top*) and a cued recall phase (*bottom*). During encoding, participants saw trial-unique object-scene pairs and indicated whether the given combination was plausible or implausible. During cued recall, participants were given either the object or the scene image as the cue and were asked to recall the paired target (scene or object image, respectively). The key conditions were (i) trials in which participants indicated they did remember the target image (“Remember” trials) and (ii) trials in which participants indicated they did not remember the target image (“Forgot” trials). Labels below denote the cue-target(memory) status of trials. O=object, S=scene, R=remember, F=forgot.

In the iEEG study, accuracy on the localiser task was on average 95% (SEM = 2%) correct (mean Reaction Time (RT) = 1.40 s, SEM = 0.21). During the cued recall task, iEEG participants indicated they remembered the target on 67% of trials (SEM = 5%). During catch trials, accuracy was 94% (SEM = 2%). RTs were faster for “Remember” trials (M = 2.59 s, SEM = 0.26) than for “Forgot” trials (M = 5.95 s, SEM = 0.40; t(10) = 8.46, P < .001). “Remember” RTs did not differ significantly for object vs. scene targets (t(10) = 0.40, P = .695).

In the scalp EEG study, participants remembered 60% (SEM = 3%) of target images. Accuracy on catch trials was 92% (SEM = 2%). RTs were faster for “Remember” trials (M = 1.61 s, SEM = 0.08) than for “Forgot” trials (M = 2.37 s, SEM = 0.17; t(19) = 5.30, P < .001). Again, RTs did not differ significantly for object vs. scene targets (t(19) = 0.73, P = .476).

### A hippocampal recall signal at ~500 ms

Our first analysis examined spectral power in the hippocampus (Figure 2A) during successful vs. unsuccessful cued recall (“Remember” vs. “Forgot”). As shown in Figure 2B, we observed an extended cluster in the gamma frequency range (60-110 Hz, 570-1730 ms, peak frequency: 85 Hz) in which “Remember” trials elicited greater power than “Forgot” trials (P_cluster_ = .007, average cluster t(10) = 3.53). The gamma effect was followed by a power decrease for “Remember” trials relative to “Forgot” trials below 30 Hz, with a distinctive peak in the alpha band (900-2600 ms, peak frequency: 9 Hz; P_cluster_ = .001, average cluster t(10) = −3.34). Hippocampal gamma and alpha power time courses are shown in Figure 2C. This finding replicates a previous report in which we found a gamma power increase followed by an alpha power decrease for successful vs. unsuccessful associative recognition memory (Staresina et al., 2016) and extends it to a cued recall paradigm.

**Figure 2.**
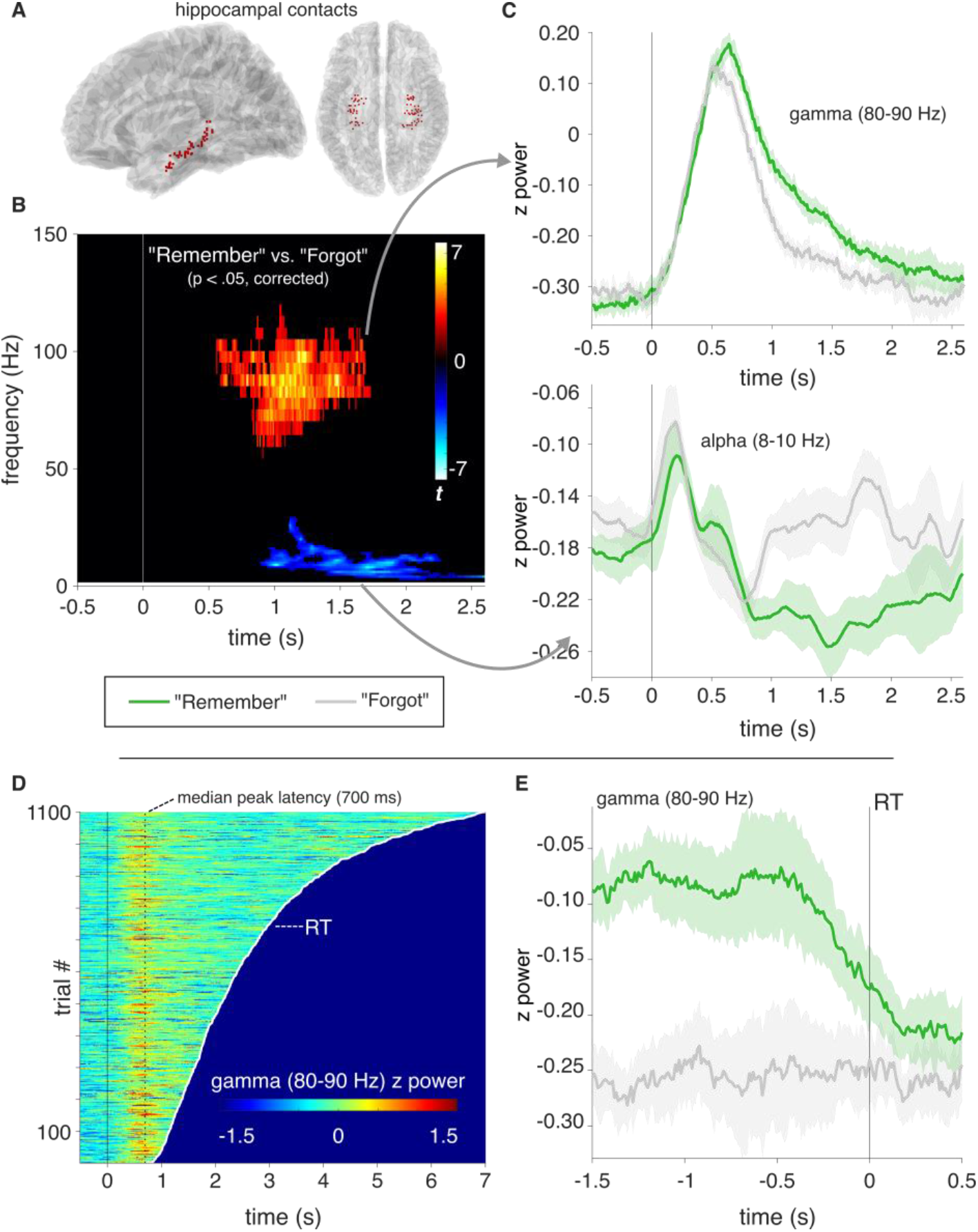
Hippocampal recall effects. **A**. Hippocampal contacts across participants shown on a normalised sagittal (*left*) and horizontal (*right*) brain template. **B**. Results from a time-frequency analysis (P < .05, corrected), contrasting “Remember” vs. “Forgot” trials and revealing a cluster in the high gamma range (60-110 Hz, peak at 85 Hz) with power increases for “Remember” trials, followed by a cluster in the alpha band (3-29 Hz, peak at 9 Hz) with power decreases for “Remember” trials. For an unthresholded map, see Figure S1 **C.** Power time courses for “Remember” (green) and “Forgot” (grey) trials in the significant gamma (*top*) and alpha (*bottom*) clusters. Lines show condition means +/− SEM of condition differences across participants. **D**. Hippocampal gamma power (80-90 Hz across time from −.5 s to reaction time (RT)) across all “Remember” trials (pooled across participants), sorted based on trial-specific RT (white line). Dashed vertical line indicates median peak latency across all trials (700 ms). **E.** Response-locked hippocampal gamma power (80-90 Hz) for “Remember” (green) and “Forgot” (grey) trials. Lines show condition means +/− SEM of condition differences across participants.

The hippocampal recall effect between ~500 and 1500 ms could emerge from two different scenarios. First, it could represent a ‘hardwired’ peak reflecting input propagation delays from visual cortex (Mormann et al., 2008), followed by sustained engagement from 500-1500 ms that accompanies the recall process. Alternatively, it could emerge from transient events (e.g., discrete bursts (van Ede et al., 2018)) occurring at different latencies across trials, with gamma peak latencies perhaps tracking reaction times (RTs). To adjudicate between these alternatives, we first plotted hippocampal gamma power (80-90 Hz) for all “Remember” trials as a function of RT. As shown in Figure 2D, this revealed a highly consistent peak at ~700 ms post cue onset, regardless of RT. Next, to test whether the hippocampal gamma effect accompanies successful recall in a sustained fashion, we plotted response-locked gamma power. As shown in Figure 2E, this revealed that the gamma effect co-terminated with a participant’s “Remember” response (significant cluster from −1500 ms to −50 ms, average t(10) = 4.28, P_cluster_ = .001). Together, these results suggest that a hippocampal recall signal sets in at ~500 ms and sustains until retrieval is complete (see Discussion).

### From perception to memory via the hippocampus

To investigate whether the hippocampal recall signal marks the switch from perceptual to mnemonic representations, we used participants’ extrahippocampal iEEG contacts (Figure 3A) to train a linear classifier on object vs. scene trials during the localiser sessions. Results confirmed high classification accuracy (Figure S2), with peak performance between 300-400 ms post stimulus onset. To capture object and scene representations during recall, a classifier was trained on the localiser data and applied to retrieval, yielding a time series of continuous object vs. scene evidence for successful and unsuccessful *object cue-scene target* and *scene cue-object target* retrieval trials (Figure 1). As shown in Figure 3B, retrieval representations were, as expected, strongly cue-driven within the first 500 ms for both “Forgot” and “Remember” trials (“Forgot” trials: 245-495 ms, P_cluster_ = .001, average cluster t(10) = 3.63; “Remember” trials: 245-445 ms, P_cluster_ = .008, average cluster t(10) = 3.86). Critically, only “Remember” trials then showed evidence for a representational switch from the cue to the target category. Specifically, for S-O(R) trials, representational patterns shifted from scene (cue) to object (target) evidence at ~500 ms and analogous for O-S(R) trials. The collapsed target evidence (i.e., the average of object evidence for S-O trials and flipped scene evidence for O-S trials) was signicantly greater than chance for “Remember” trials from 720-900 ms (P_cluster_ = .023, average cluster t(10) = 2.57). Lastly, this collapsed target evidence was significantly greater for “Remember” trials than for “Forgot” trials from 555-895 ms (P_cluster_ = .001, average cluster t(10) = 3.15).

**Figure 3.**
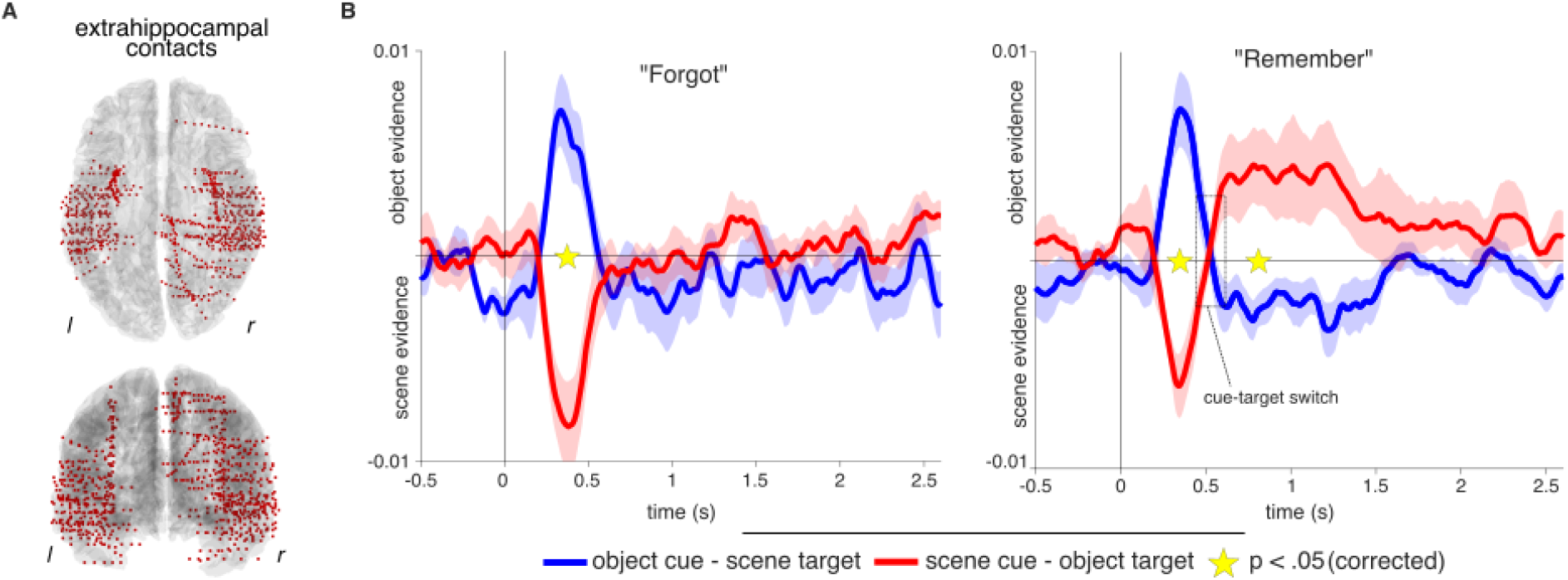
Switch from cue to target representations. **A**. Extra-hippocampal contacts across participants shown on a normalised horizontal (*top*) and coronal (*bottom*) brain template. **B**. Object and scene evidence, based on a classifier trained on a ‘localiser’ session, for O-S (blue) and S-O (red) trials, separately for “Forgot” (*left*) and “Remember” (*right*) trials. Within the first 500 ms, classifier evidence reflects the cue category. Only for “Remember” trials, classifier evidence then switches (dashed rectangle) to reflect the recalled target category.

Inspection of gamma power (Figure 2C) and classification time courses (Figure 3B) raises the intriguing possibility that the hippocampal gamma power increase during “Remember” trials at ~500 ms marks the moment at which the brain switches from cue to target representations. Figure 4A shows the hippocampal gamma recall effect (80-90 Hz, “Remember” minus “Forgot”) superimposed on the classification effect, suggesting that these two effects indeed evolve in tandem. To assess this notion more directly, we repeated the classification analysis for “Remember” trials, but realigned each trial to its hippocampal gamma power peak. Figure 4B (left) illustrates the switch from cue to target evidence around the hippocampal gamma peak. Note that the hippocampal recall effect sets in ~100 ms prior to the actual gamma peak (Figure 2, gamma effect onset: 570 ms, gamma peak for “Remember”: 700 ms), consistent with the notion that the representational cue-target switch starts as hippocampal gamma power distinguishes “Remember” from “Forgot” trials. To quantify the representational switch around hippocampal gamma peaks statistically, we averaged classifier evidence for O-S and S-O “Remember” trials across a 500 ms pre- and a 500 ms post-hippocampal gamma peak window (Figure 4B, right) and conducted a repeated-measures ANOVA including the factors Time Window (pre, post) and Trial Type (O-S, S-O). Results revealed a significant interaction (F(1,20) = 16.76, P = .002) in the absence of any main effect (both P > .28). For O-S trials, classifier evidence changed significantly from object (cue) to scene (target) evidence (t(10) = 2.73, P = .021). Likewise, for S-O trials, classifier evidence changed significantly from scene (cue) to object (target) evidence (t(10) = 4.14, P = .002). Finally, the same interaction emerged when comparing classifier evidence in a cue-locked vs. hippocampal-peak-locked fashion (F(1,20) = 15.62, P = .003, again using 500 ms windows). Together, these results corroborate the notion that hippocampal engagement during “Remember” trials marks the moment of an extrahippocampal switch from perceptual cue to mnemonic target representations.

**Figure 4.**
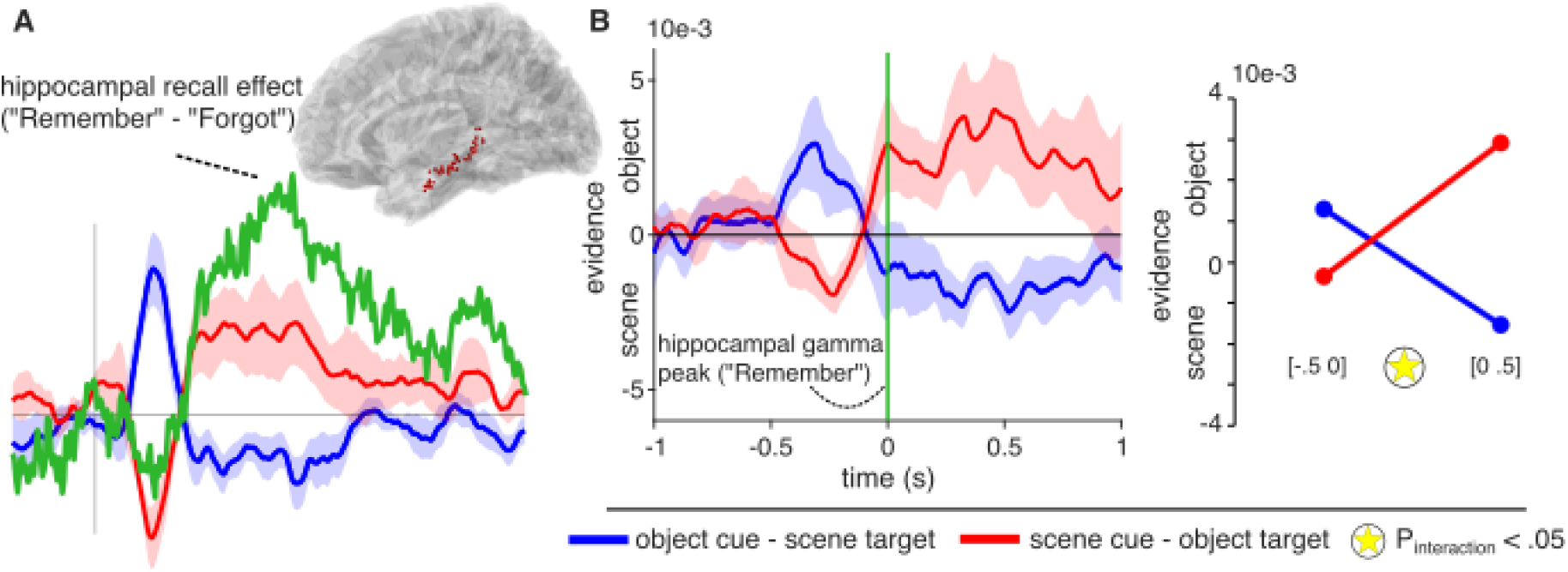
A hippocampal switchboard function. **A**. Overlay of the hippocampal recall effect (“Remember” minus “Forgot” gamma power, cf. Figure 2C, a.u.) onto the extrahippocampal switch from cue to target representation for “Remember” trials (cf. Figure 3B). Inset shows hippocampal contacts across patients. **B**. *Left*: Object and scene evidence for “Remember” trials as in Figure 3B, but realigned to trial-by-trial hippocampal gamma peaks (time 0, vertical green line). Note the cue evidence (scene for S-O trials, object for O-S trials) before the hippocampal gamma peak vis a vis target evidence (object for S-O trials, scene for O-S trials) after the hippocampal gamma peak. *Right*: Illustration of the significant Time Window x Trial Type interaction around the gamma peak. [−.5 0] and [0 .5] denote the 500 ms time windows across which the classifier data were averaged.

**Figure 4.**
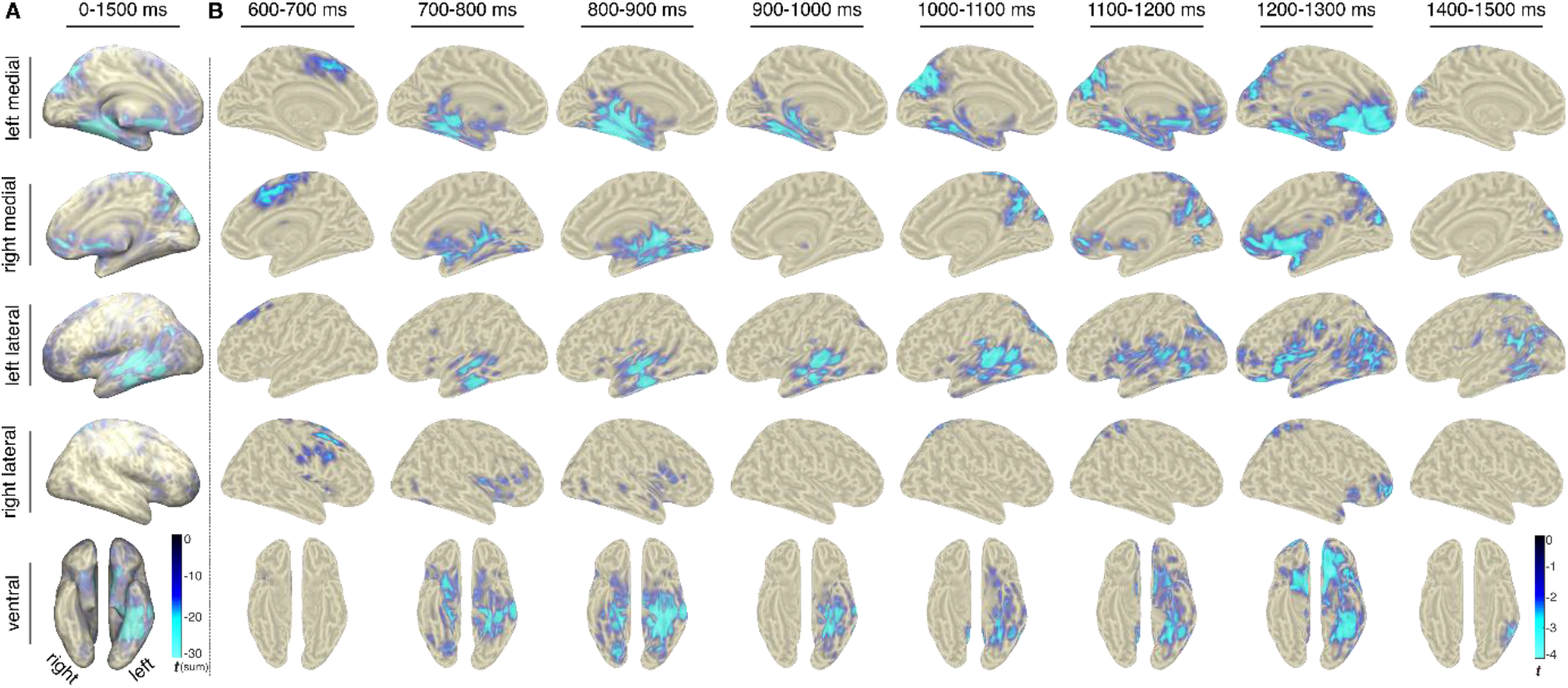
Brain-wide recall dynamics. Results from source-localised EEG alpha (8-10 Hz) power, comparing “Remember” vs. “Forgot” trials. **A**. Voxel x time results (P_cluster_ < .05), revealing recall effects across the core retrieval network (including hippocampus, PPC, LTC and vmPFC). Colour reflects sum of significant t values across time. **B**. Time-resolved results (proceeding in 100 ms steps), showing only time windows with significant effects (each map thresholded at P_cluster_ < .05, no significant effects emerged from 1300-1400 ms).

### Recall signals across the cortical retrieval network

Results from our iEEG sample showed that ~500 ms after cue onset, a hippocampal signal distinguishes successful from unsuccessful cued recall and marks the switch from cue- to target representations. However, hippocampal engagement alone is unlikely to be sufficient for full-blown memory recall. To elucidate the cortical dynamics associated with cued recall, we conducted the same experiment in a sample of 20 healthy participants using high-density scalp EEG.

A first sensor x frequency x time comparison of “Remember” vs. “Forgot” revealed an extended cluster of relative power decreases for “Remember” trials (Figure S3), with a peak effect at again 9 Hz (as for the iEEG study, cf. Figure 2B). We next projected the sensor EEG data into source space (see Methods), extracted alpha power (8-10 Hz) in the resulting virtual voxels and examined how the recall signal (alpha power decrease for “Remember” compared to “Forgot” trials) evolves across time. Performing the analysis across the entire 0-1500 ms interval revealed significant recall effects across the CRN, including MTL, posterior parietal cortex (PPC), lateral temporal cortex (LTC) and ventromedial prefrontal cortex (vmPFC; Figure 4A). This result replicates and extends a previous study in which we used source-localised Magnetoencephalography (MEG) alpha power to reveal recall effects across the CRN (Martín-Buro et al., 2020).

Critically, we next performed the same analysis in a time-resolved fashion, progressing from 0 to 1500 ms in 100 ms steps, each step averaging alpha power across a 100 ms time window. Each comparison was again cluster-corrected for multiple comparisons across virtual voxels. As shown in Figure 4B, the first time window to show a significant recall effect occurred from 600-700 ms and encompassed dorsomedial prefrontal cortex (dmPFC)/anterior cingulate cortex (ACC). Next, a significant effect was seen from 700-1000 ms in the MTL. It is worth pointing out that this is also the time window in which our intracranial data showed an alpha power decrease for “Remember” trials in the hippocampus (Figure 2B), providing a link between the two datasets and imaging modalities. From ~900-1300 ms the recall effect encompassed medial and lateral PPC, followed by a recall effect in vmPFC from ~1100 – 1300 ms.

To quantify these latency differences statistically, we extracted alpha power (8-10 Hz) time courses in voxels showing significant effects in the initial voxel × time contrast (Figure 4A) and overlapping with anatomically defined regions of interest for dmPFC, hippocampus, medial PPC and vmPFC. The alpha power effect (“Remember” vs. “Forgot”) was then binned into ten 100 ms segments from 500-1500 ms and subjected to a repeated-measures ANOVA (Greenhouse-Geisser corrected) including the factors Region and Time. Results confirmed a significant Region x Time interaction (F(6.46, 122.74) = 2.59, P = .019) in the absence of any main effect (both P > .380). Time courses and effect latencies are illustrated in Figure S4. Together, these results reveal a hitherto unknown sequence of recall signals within the CRN, starting in dmPVC/ACC and then proceeding from hippocampus via PPC to vmPFC.

## Discussion

Our study elucidates the role of the hippocampus as a switchboard from perception to memory and unveils the ensuing cortical dynamics supporting the recall process. Using a simple and robust cued recall paradigm (Figure 1), iEEG recordings first revealed a hippocampal signal in the high gamma range (60-110 Hz) distinguishing between successful and unsuccessful recall from 500 ms onwards (Figure 2B). This gamma effect was followed by a relative power decrease for successful recall in the alpha band. Using multivariate pattern analysis (MVPA), we observed – for successful recall only – a representational switch from the cue stimulus category (< 500 ms) to the target stimulus category (> 500 ms; Figure 3B). Time-locking the MVPA to hippocampal gamma peaks showed cue evidence before and target evidence after these peaks, indicating that the hippocampal gamma increase marks the moment at which brain states shift from perceptual to mnemonic representations. Moving beyond the hippocampus with high-density scalp EEG, we first established engagement of the cortical retrieval network (CRN) during successful recall, including posterior parietal cortex (PPC) and ventromedial prefrontal cortex (vmPFC) (Figure 4A). Critically, using time-resolved alpha power in source space, we found a particular recall cascade across the CRN: Starting at ~700 ms in the MTL, successful recall subsequently entailed PPC at ~900 ms, followed by vmPFC at ~1100 ms.

### The hippocampus as the switchboard from perceptual cues to mnemonic targets

Our data provide strong empirical evidence for the long-held notion that the hippocampus orchestrates cortical pattern completion (Marr, 1971; Norman and O’reilly, 2003; Teyler and DiScenna, 1986). Owing to modality-specific methodological limitations across species, such empirical evidence has been challenging to obtain. That is, fMRI lacks the temporal resolution to pinpoint a hippocampal signal preceding target reinstatement, although recent analytical advances have yielded some progress in resolving fine-grained memory dynamics with fMRI (Staresina et al., 2013; Wittkuhn and Schuck, 2021). Scalp EEG and MEG, combined with advanced source reconstruction methods (Gross, 2019; Michel et al., 2004), in principle provide adequate levels of spatial and temporal precision to uncover whole-brain memory dynamics (Bergström et al., 2013; Martín-Buro et al., 2020). However, ambiguities remain when interpreting activation in deeper sources such as the medial temporal lobe (MTL), at least without converging evidence from other imaging modalities. Optogenetic studies in mice have shown that experimental activation of hippocampal cell assemblies elicits contextual fear behaviour (Liu et al., 2012) and that silencing hippocampal cells abolishes reinstatement of memory representations in cortical structures such as entorhinal cortex, perirhinal cortex and retrosplenial cortex (Tanaka et al., 2014). However, it remains open to what extent contextual fear conditioning captures the intricacies of episodic memory recall in humans. Moreover, these studies remain agnostic about the fast temporal relationship between hippocampal and cortical engagement during episodic memory recall.

The hippocampal recall effect reported here (Figure 2) unifies and extends a series of recent human electrophysiological results. Specifically, a hippocampal recall signal starting at ~500 ms after onset of a retrieval cue has been reported for evoked field potentials (Staresina et al., 2012a), high gamma power (Staresina et al., 2016) and single neuron firing rates (Staresina et al., 2019), attesting to the complementary nature of these electrophysiological signals (Buzsáki et al., 2012). Figure 2 moreover illustrates the sustained nature of the hippocampal recall effect, extending from ~500-1500 ms post cue onset. We have interpreted the 500 ms onset as reflecting conduction delays from sensory regions to the hippocampus (Mormann et al., 2008) and the ensuing ~1 s period as reflecting recurrent hippocampal-cortical interactions in service of memory retrieval (Staresina and Wimber, 2019). However, this pattern could also emerge from transient bursts (Vaz et al., 2019; Vaz et al., 2020) igniting the recall process at different latencies across trials, perhaps tracking trial-specific response times (RTs). As shown in Figure 2D (left), our data point to a highly consistent gamma peak occurring at ~700 ms post cue onset irrespective of RT, corroborating the notion that it reflects a relatively ‘hard-wired’ delay at which a hippocampal recall signal sets in, at least in the experimental context of our cued recall paradigm. This gamma peak latency agrees with previous studies examining single neuron firing latencies in memory-selective hippocampal neurons (Rutishauser et al., 2015; Staresina et al., 2019) and event-related potential (ERP) recordings from the hippocampus (Smith et al., 1986). Importantly though, we also found that hippocampal gamma remains sustained until a memory response is given (Figure 2D, right), suggesting that hippocampal engagement accompanies extrahippocampal reinstatement processes throughout recall. It deserves mention that the consistent onset notwithstanding, the sustained recall signal may well include discrete gamma bursts/ripples occurring at different latencies across trials (van Ede et al., 2018).

While we here focused on target reinstatement in extrahippocampal sites, theoretical models implicate a prior pattern completion process within the hippocampus to retrieve the ‘index’ of the target representation (Teyler and DiScenna, 1986; Teyler and Rudy, 2007). In the current paradigm, it is challenging to disentangle whether any similarity between a given retrieval trial and its encoding counterpart would reflect such pattern completion processes or the perceptual match of the cue image with the encoding display (Figure 1). That said, in a previous study we found that an intra-hippocampal pattern completion process commenced ~500 ms after cue onset (Staresina et al., 2016) and directly correlated with high gamma power increases. Together, these data suggest that at ~500 ms, a cue representation reaches the hippocampus and induces an intrahippocampal pattern completion process. If successful (reflected in increased high gamma power), this ignites sustained reinstatement of the episodic target representation in cortex (Staresina and Wimber, 2019). The subsequent decrease in hippocampal alpha power might then reflect increased levels of information processing, as postulated and shown for cortical information processing (Griffiths et al., 2019; Hanslmayr et al., 2016). As elaborated below, this alpha power decrease subsequently tracks the dynamic recall signal throughout the cortical retrieval network.

### Temporal dynamics in the cortical retrieval network

The process of reinstating a full-blown episodic memory and deploying adaptive behaviour most likely relies on intricate interactions across multiple cortical areas beyond the hippocampus (Olsen and Robin, 2020; Ritchey and Cooper, 2020). Apart from content-specific areas involved in reinstatement (Ritchey et al., 2013; Staresina et al., 2012b; Wheeler and Buckner, 2004), recent fMRI research has revealed a cortical brain network consistently emerging during successful recall (Rugg and Vilberg, 2013). This network includes medial and lateral parietal cortex (PPC) and ventromedial prefrontal cortex (vmPFC), all of which are densely connected with the hippocampus (Aggleton, 2012; Ferguson et al., 2019). What is still unresolved, however, is the particular role each of the different cortical retrieval network (CRN) nodes plays (Ritchey and Cooper, 2020).

Capitalising on the temporal resolution of EEG, we found that within the CRN, a recall effect (alpha power decreases) first emerged in the MTL (including hippocampus, see also Figure S4), followed by medial PPC and lastly in vmPFC (Figure 4). Involvement of the hippocampus in this recall task is corroborated by our intracranial data revealing a hippocampal alpha power effect spanning the same time and frequency window (Figure 2) as well as by a previous fMRI study using the same paradigm (Staresina et al., 2013). This result adds to recent evidence emphasising the feasibility of using source-localised M/EEG recordings to examine hippocampal memory processes (Martín-Buro et al., 2020; Pizzo et al., 2019; Pu et al., 2018; Ruzich et al., 2019).

In any case, the functional significance of the MTL-PPC-vmPFC trajectory is unknown at present. There is ongoing debate about the role of different PPC regions (e.g., posterior midline, superior parietal lobule, inferior parietal lobule) in episodic retrieval (Cabeza et al., 2008; Wagner et al., 2005), but one prevalent view is that involvement of PPC regions scale with the amount of mnemonic evidence (Wagner et al., 2005). Likewise, the functional parcellation and the specific role of medial PFC in memory retrieval is still poorly understood (De La Vega et al., 2016), although there is consensus about a role of prefrontal areas in higher order information integration and action planning (Gilbert et al., 2006; Rushworth et al., 2004). One tentative scenario could thus be that PPC serves as an ‘episodic buffer’ (Vilberg and Rugg, 2008), accumulating episodic details that are reinstated in content-specific areas through hippocampal pattern completion. Ventromedial PFC might then integrate this mnemonic evidence with the current task set and initiate goal-directed behaviour. On that note, it deserves mention that the MTL alpha power effect was preceded by a recall effect in dorsomedial prefrontal cortex (dmPFC)/anterior cingulate cortex (ACC) (Figure 4). This could reflect a general top-down/control mechanism influencing the hippocampus and/or may reflect the role of ACC specifically in conflict resolution (Botvinick et al., 2004), given that the external/perceptual cue image needed to be overridden in favour of the internal/mnemonic target image in our paradigm.

Finally, the temporal dynamics of our findings are consistent with a role of the hippocampus - perhaps under top-down control of dmPFC - in triggering the switch from perception to memory and the associated recall cascade in the CRN. However, any conclusive evidence for a causal role would require an interventionist approach. This may be afforded by future work employing intracranial electrical stimulation of the hippocampus and/or perturbation of particular CRN nodes at specific times via transcranial magnetic stimulation (TMS).

## Materials and methods

### Participants

For the intracranial EEG study, 10 patients from the Queen Elizabeth Hospital in Birmingham (UK) and one patient from La Paz University Hospital in Madrid (Spain), all suffering from medically intractable epilepsy, volunteered (6 male, 5 female, aged 24-53, M = 34.45). Additional patient characteristics are listed in Table S1. Ethical approvals were granted by the National Research Ethics Service UK (code 15/EM/0182) and by the Clinical Research Ethics Committee at La Paz University Hospital Madrid (code IP-2401), respectively.

Twenty healthy, right-handed participants (12 male, 8 female) with normal or corrected-to-normal vision volunteered in the EEG experiment. They were aged 20-33 years (M = 25.01). An additional six participants had been rejected from analysis due to noisy EEG data (n=2), inconsistent Polhemus data (n=2), or poor memory performance (< 40% ‘Remember’ trials, n=2). All participants were fluent English speakers. Participants gave written informed consent and received course credits or financial remuneration. Ethical approval was granted by the University of Birmingham Research Ethics Committee (ERN_14-1379). In additional to functional recordings, structural MRIs were acquired for 15 participants.

### EEG experimental procedure

The stimulus material consisted of 712 colour images sized 200 x 200 pixels, half depicting objects and half depicting scenes. It was based on a set of images used in previous studies (Konkle et al., 2010; Staresina et al., 2013) supplemented with additional images obtained via a Google search that matched the main image set in style. Participants received written and verbal instructions.

Before and after the main experiment, participants performed ‘localiser’ runs. In each run, participants completed 10 practice trials (5 objects, 5 scenes) that were not recorded followed by 100 unique images (50 objects, 50 scenes) presented in the centre of the screen. Each trial started with a fixation cross presented for 1.5 ± 0.1 s. Subsequently, an object or scene image was superimposed on the fixation cross. Participants had to press a button to indicate whether the image depicts an object or a scene. After 1 s, a legend appeared at the bottom of the screen reminding participants of the assignment between left/right buttons and object/scene. To avoid contamination of the classifier by response mapping, this assignment was flipped in the second localiser run (initial assignment counterbalanced across participants). The trial terminated after a button press, although the image was shown for a minimum of 2 s and a maximum of 10 s. The localiser was included in the EEG study to match the iEEG paradigm, but data are not used in the present manuscript.

The main experiment consisted of 8 runs following the paradigm used in (Staresina et al., 2013). Each run was split into 4 blocks: a pre-encoding delay block, an encoding block, a post-encoding delay block, and a retrieval block. Before and after each block, a progress bar was displayed for 6 s, alerting participants to the impending start of the next block. During delay blocks, random numbers between 0 and 100 were shown, and participants pressed the left key for even numbers and the right key for odd numbers. This phase was self-paced, with a new number appearing immediately after a button press. Participants were encouraged to perform the task as fast as possible while maintaining high performance. Each delay block lasted 3 min.

Each encoding block consisted of 32 trials. Each trial started with a fixation cross presented for 1.5 ± 0.1 s. Subsequently, a unique, randomly chosen object-scene pair was shown. During 16 randomly assigned trials, the object appeared left of the centre and the scene appeared right, with the opposite arrangement for the other 16 trials. The object-scene pair remained on the screen until a button was pressed, but it was displayed for a minimum of 2.5 s and a maximum of 4 s. Participants used their right hand and indicated with the index finger that the object-scene pair was “plausible”, i.e., likely to appear in real life or nature, or used their middle finger to indicate that it was “implausible”.

Each retrieval block comprised 32 trials. Each trial commenced again with a fixation cross 1.5 ± 0.1 s. Subsequently, a cue was shown in the centre of the screen, either an object or a scene taken from the previous encoding block. The object-scene pair remained on the screen until a button was pressed, but it was displayed for a minimum of 2.5 s and a maximum of 6 s. Participants were asked to indicate whether they “remember” (index finger) or “forgot” (middle finger) the corresponding paired associate. Half of the cues were objects, the other half scenes. Across the 32 trials, each cue type (object or scene) was presented in mini-blocks of eight consecutive trials alternating between 8 object cues (O) and 8 scene cues (S), i.e., O-S-O-S or S-O-S-O. Participants were instructed to only press “Remember” when their memory was vivid enough to give a detailed description of the associate. To ensure that this is indeed the case, in 20% of the cases “Remember” responses were followed by the instruction to enter a description of the target associate using the computer keyboard (‘catch-trials’). The experiment was programmed with Psychophysics Toolbox Version 3 (Brainard, 1997).

### iEEG experimental procedure

For the iEEG study, the procedure was largely similar, with a few modifications. The stimulus pool for the memory portion consisted of 192 objects and 192 scenes (drawn from the same pool as described above). To accommodate different levels of cognitive capacity across epilepsy patients, we prepared three versions of the experiment, varying in the duration of each run. In Level 1, an encoding/retrieval block consisted of 8 trials, resulting in a total of 24 runs including a pre/post-encoding delay of 30 s. In Level 2, there were 12 runs with 16 trials per encoding/retrieval block and 60 s delay periods. In Level 3, there were 6 runs with 32 trials per encoding/retrieval block and 120 s delay periods. Which version was used depended on performance on a short practice run at difficulty Level 1. Two patients performed the task at Level 1, four at Level 2 and the remaining five patients at Level 3. In terms of stimulus timing, responses were self-paced, but images remained on the screen for a minimum of 2 s and a maximum of 10 s (at which point the response was coded as ‘invalid’ and included in the ‘Forgot’ condition). Instead of typing in responses during the ~20% catch trials, patients verbally described the paired associate and responses were transcribed by the experimenter. In order not to overtax patients, runs were spread across 1-3 experimental sessions, with an effort to keep sessions close in time. Again, object/scene localiser runs were conducted before and after each memory session. The same set of 50 object and 50 scene images was used repeatedly, images and response legend remained on the screen for a minimum of 2 s and a maximum of 10 s, and no switch of response finger assignment was introduced.

### iEEG acquisition and preprocessing

Intracranial EEG (iEEG) was recorded for pre-surgical epilepsy diagnosis using laterally implanted depth electrodes. Electrode shafts contained 5-15 contacts. Data were digitised at 512 Hz (n=1) or 1024 Hz (n=10). Intra- and extrahippocampal contacts were identified based on the post-implantation structural MRI. Contacts with hardware artifacts were discarded based on visual inspection (average of 5% across patients). The numbers of contacts per patient included in the analyses are listed in Table S1. For hippocampal contacts, data were locally re-referenced to a white-matter contact on the same electrode. For extrahippocampal contacts, a Common Median Reference including all contacts was used to re-reference the data.

### EEG acquisition and preprocessing

Electroencephalogram (EEG) was recorded with 128 sintered Ag/AgCl active electrodes and a BioSemi Active-Two amplifier. The signal was digitized at a rate of 1024 Hz on a second computer via ActiView recording software (BioSemi, Amsterdam, Netherlands). Electrode positions and headshape were measured using a Polhemus FASTRAK device (Colchester, VT, USA) in conjunction with Brainstorm (Tadel et al., 2011). All EEG data processing was performed in MATLAB using FieldTrip (Oostenveld et al., 2011). Data were downsampled to 256 Hz, high-pass filtered at 0.1 Hz using a windowed sinc FIR filter and low-pass filtered at 100 Hz using a Butterworth IIR filter. Furthermore, a band-stop filter was applied at 50 Hz to remove line noise. Retrieval data were then segmented into epochs, starting at −1 s and ending at the time of the button press +1 s, or at 6 s post-stimulus, whichever was shorter. Noisy EEG channels were identified by visual inspection and discarded. Subsequently, Infomax ICA (Bell and Sejnowski, 1995) was used to clean the data. To this end, all epochs were manually inspected and artifact trials containing muscle artifacts or mechanical artifacts (≈10%) were discarded. The resultant data were high-pass filtered above 1 Hz and ICA was applied. Using visual inspection of the spatial patterns, time-series and power spectra, ICA components associated with eye blinks, eye movements, and EMG were rejected from the original data (prior to manual artifact rejection and high-pass filtering). Next, discarded EEG channels that were interpolated using a weighted average of the neighbouring channels. Finally, the data were re-referenced using a Common Average Reference (CAR).

### EEG source modelling

Individual structural MRIs were segmented into grey matter, white matter, cerebrospinal fluid, skull, and scalp compartments. For 5 participants, individual MRIs were not available and the standard MNI template was used instead. As geometric model of the head, a hexahedral mesh with a shift of 0.3 was used. The FieldTrip-SimBio pipeline (Vorwerk et al., 2018) was used with tissue conductivities of 0.33, 0.14, 1.79, 0.01, and 0.43 in order to create a Finite Element Method (FEM) volume conduction model. To co-register the MRI with the Polhemus coordinates of the electrodes, the fiducials (nasion, LPA, RPA) were manually identified in each MRI. A source grid model with 10 mm spacing and 3294 grid points was defined in MNI space and mapped onto each participant’s MRI. This ensured that a given grid point corresponded to the same anatomical location across participants.

The data were projected into source space using linearly constrained minimum-variance (LCMV) beamformers (Van Veen et al., 1997). Retrieval trials were band-pass filtered in the 2 – 30 Hz band and trial-wise covariance matrices were averaged and regularized.

For region-of-interest based analyses in source space, we used anatomical masks provided by the AAL atlas (Tzourio-Mazoyer et al., 2002). The following bilateral masks were included: dmPFC: ‘Cingulum_Ant’, ‘Cingulum_Mid’, ‘Supp_Motor_Area’; Hippocampus: ‘Hippocampus’; PPC: ‘Precuneus’, ‘Cingulum_Post’; vmPFC: ‘Frontal_Med_Orb’, ‘Frontal_Sup_Orb’, ‘Rectus’.

### Time-frequency analysis

For both iEEG and EEG, short-time Fourier analysis of the retrieval data was performed using FieldTrip with sliding time windows in 10-ms (iEEG) or 25-ms (EEG) steps. For a lower frequency range (2–29 Hz iEEG, 2-48 EEG, 1-Hz steps), the window length was set to five cycles of a given frequency and the windowed data segments were multiplied with a Hanning taper. For calculation of EEG alpha power (8-10 Hz) in source space, 50-ms time steps were used. For iEEG hippocampal gamma power (30– 150 Hz, 5-Hz steps, iEEG only), we applied multitapering using a fixed window length of 400 ms and seven orthogonal Slepian tapers. Power values for each frequency were normalised via z-transformation across all trials, including a 500 ms pre-stimulus baseline interval. To remove outliers, the 10% most extreme power values across trials were discarded within each channel and for each time/frequency bin prior to creating condition-specific averages. Analyses were restricted to 2.6 s post cue onset for iEEG data and to 1.5 s post cue onset for EEG data, as this marked the respective average response time for “Remember” trials across participants (leaving a 100 ms buffer in the EEG data to avoid contamination by the motor response).

### Multivariate analysis

Multivariate classification of iEEG data was performed using MVPA-Light (Treder, 2020). For all multivariate analyses, a linear discriminant analysis (LDA) was used as classifier (Fisher, 1936). The classifier was trained on localiser data to discriminate between objects and scenes. When applied to the retrieval data, the classifier produces a decision value (dval) that represents the signed distance to the hyperplane. Here, a positive dval is evidence for an activation pattern associated with an object, whereas a negative dval is associated with a scene.

For both localiser and retrieval data, z-scoring was applied across trials for each time point separately to normalise channel variances and remove baseline shifts. Z-scoring was first done across trials within each run in order to account for signal changes across time. The runs were then concatenated and jointly normalised using another z-scoring operation. Non-hippocampal contacts (36-128 depending on participant) served as features. To quantify whether object and scene cues can be differentiated, LDA was trained and tested on the localiser data (Figure S1, 100 ms temporal smoothing applied). Data were split into training and test sets using 5-fold cross-validation (Lemm et al., 2011) and Area Under the ROC Curve (AUC) was used as metric. The analysis was repeated five times with random folds in each iteration and results were averaged. To investigate the occurrence of object/scene representations in the retrieval phase (Figure 3), **Error! Bookmark not defined.** used a transfer learning approach wherein the classifier was first trained on the localiser data averaged in the 300-400 ms window that contained the peak performance. This classifier was then tested for every time point in the retrieval phase (also applying 100 ms temporal smoothing).

To investigate how the timing of the switch from cue to target representation relates to hippocampal gamma power, we realigned each retrieval trial to its respective gamma peak. To this end, time-frequency data were normalised in each frequency band as described above and a single power time series was created by averaging z power across hippocampal contacts within the 60-110 Hz range. Again, prior to averaging across frequencies and hippocampal contacts, a trimmed mean was used wherein power values above the 90-th percentile (across trials) were discarded. We then identified local maxima in the 0.2 – 1.2 s interval cantered on the median latency of gamma peaks across participants for “Remember” trials (0.7 s). If one or more discrete gamma peaks were found in this interval, the respective time axis was realigned to the highest peak. Transfer classification was then repeated on the realigned data and dvals were calculated.

### Statistics

For behavioural analyses, reaction times (RTs) within participants were summarised by calculating the median in order to mitigate the effect of outliers. At the group level, arithmetic mean (M) and standard error of the mean (SEM) are reported. Paired-samples t-tests were used to compare RTs in “Remember” and “Forgot” trials, and for object and scene cues (in “Remember” trials). Unless stated otherwise, FieldTrip’s cluster permutation test (Maris and Oostenveld, 2007) was used to account for multiple comparisons for all time-frequency and classification analyses, both in sensor space and in source space. A paired-samples t-test with a threshold of P < .05 was used to define initial clusters. Maxsum (sum of all t-values in cluster) served as cluster statistic and Monte Carlo simulations were used to calculate the cluster p-value (alpha = .05, two-tailed) under the permutation distribution. Analyses were performed at the group level.

## Resource Availability

Data and analysis code will be made publicly available on OSF (https://osf.io/) upon publication.

## Acknowledgements

This work was supported by a Wellcome Trust/Royal Society Sir Henry Dale Fellowship (107672/Z/15/Z) to B.P.S., a European Research Council Starting Grant (ERC-2017-StG 759432) to I.C., a European Research Council Consolidator Grant (647954) to S.H., a grant from the Wolfson Foundation and Royal Society to S.H., a grant from the Economic Social Sciences Research Council (ES/R010072/1) to S.H. and a European Research Council Starting Grant (ERC-STG-2016-715714) to M.W.

## Author Contributions

Conceptualization, M.T., B.P.S.; Methodology, M.T., B.P.S.; Software, M.T., I.C., S.M.; Formal Analysis, M.T., B.P.S.; Investigation, M.T., F.R., M.C.M.-B., B.P.S.; Resources, F.C.B., A.U.-C., V.S., R.C., S.H.; Data Curation, M.C.M.-B., D.T.R.; Writing – Original Draft, M.T., B.P.S.; Writing – Review & Editing, M.T., I.C., M.W., S.H., B.P.S.; Visualization, M.T., B.P.S.; Supervision, B.P.S.; Project Administration, B.P.S.; Funding Acquisition, S.H., B.P.S.

## Supplemental Material

**Table S1.**
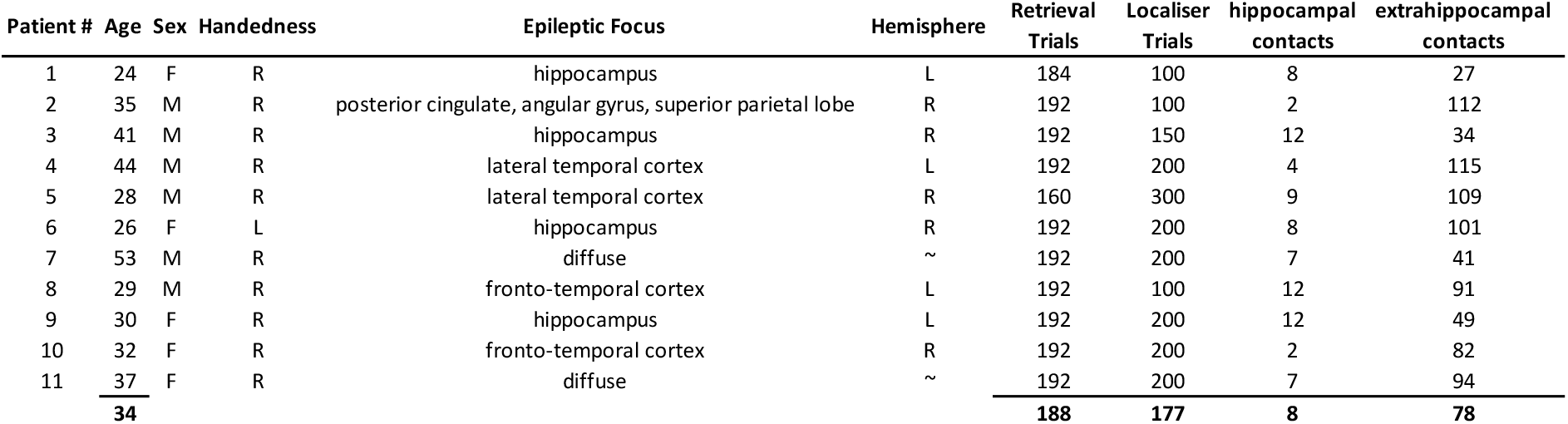
iEEG patient characteristics. Bold numbers denote group averages. ‘Diffuse’ epileptic focus indicates that invasive monitoring did not yield a clear focus.

**Figure S1.**
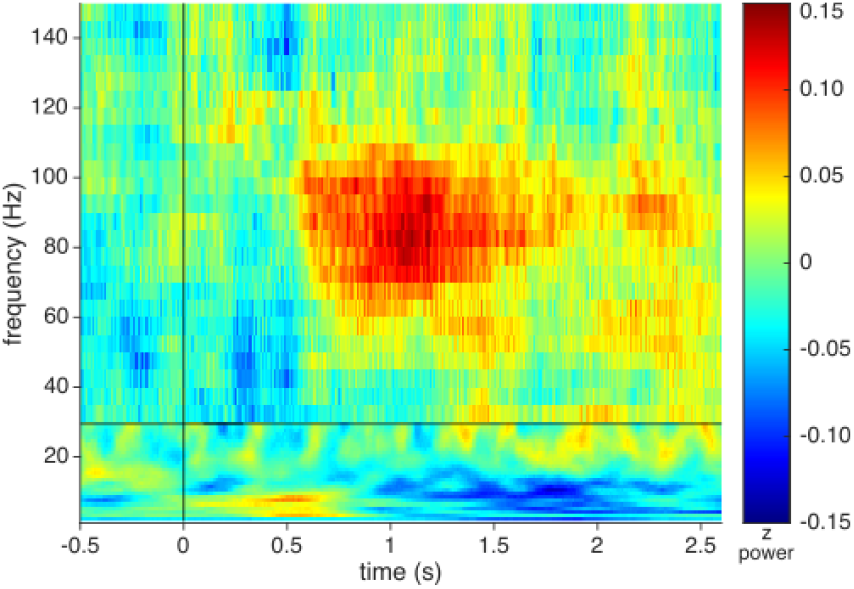
Unthresholded hippocampal time-frequency map. Colours depict the difference in z power for “Remember” vs. “Forgot” trials, averaged across participants. Horizontal black line at 30 Hz indicates different settings for deriving power for lower vs. higher frequencies.

**Figure S2.**
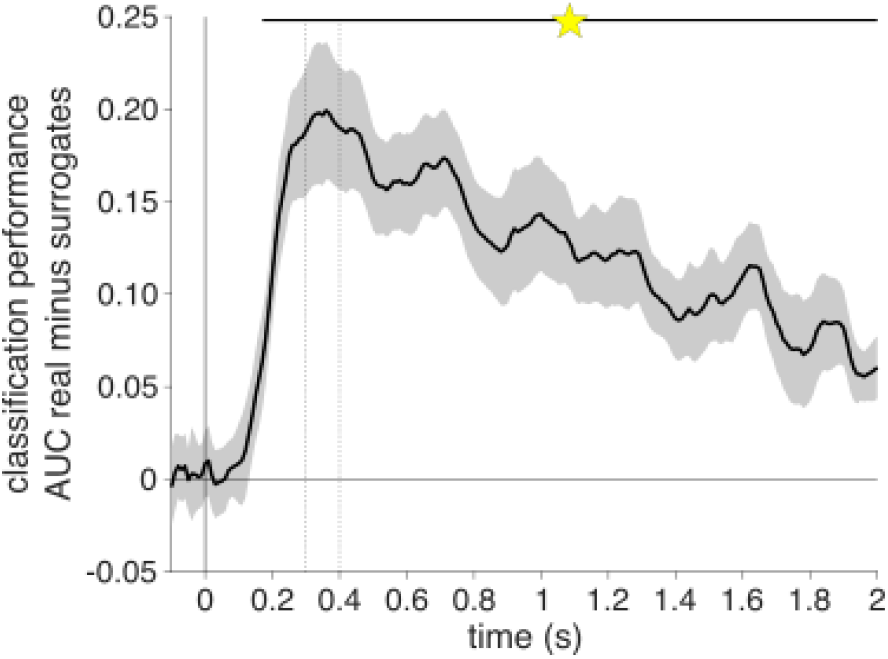
iEEG object vs. scene decoding. Results show mean +/−SEM of cross-validated LDA results across participants, revealing significant above-chance classification (relative to label-shuffled surrogates) from ~200 ms onwards, with peak performance from 300–400 ms (dashed vertical lines). Horizontal black line indicates statistical significance using cluster-based correction for multiple comparisons across time (P_cluster_ < .001)

**Figure S3.**
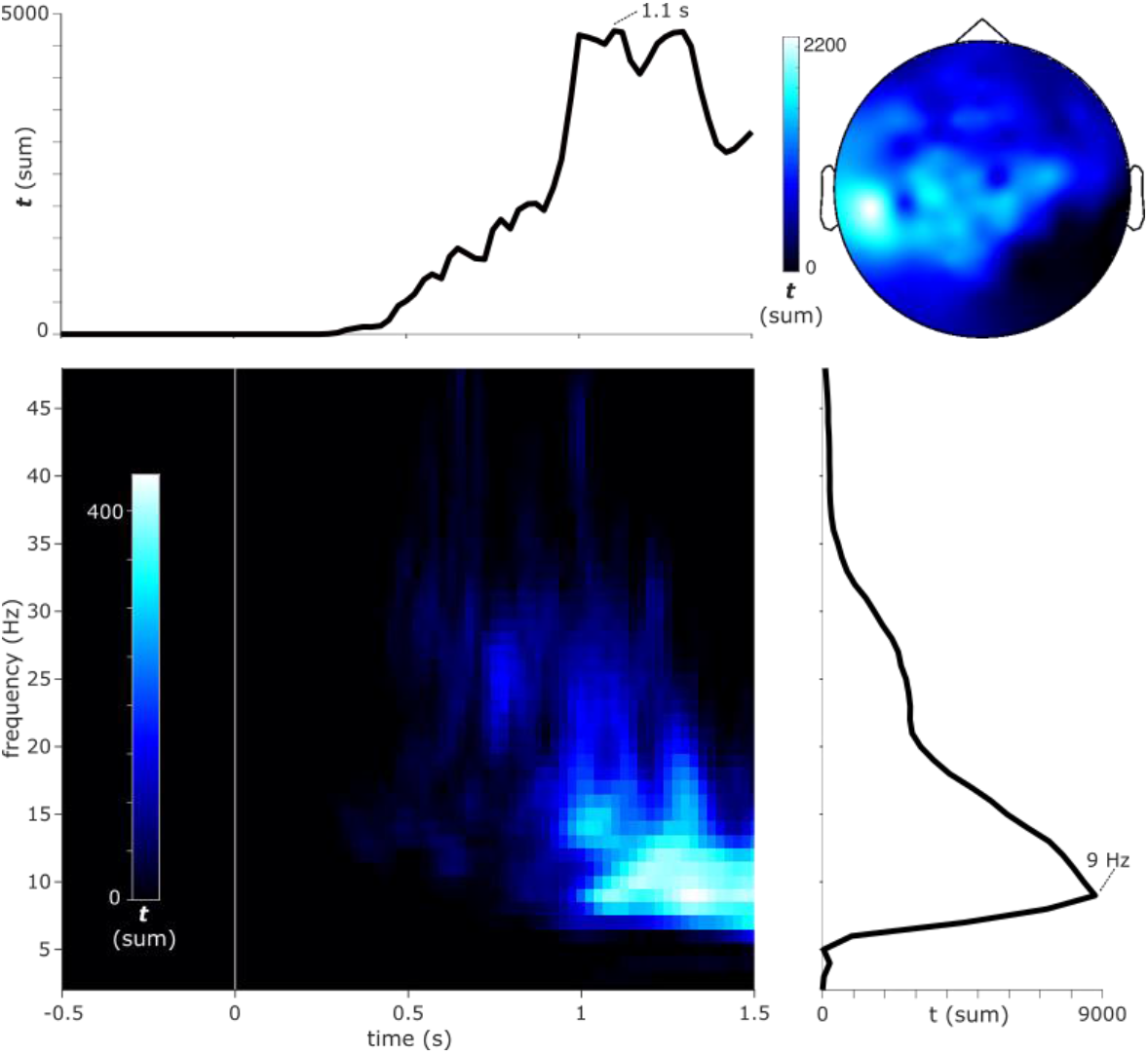
Extended EEG results of time x frequency x sensor comparison for “Remember” vs. “Forgot” trials, including a time range from 0–1.5 s, a frequency range from 2–48 Hz and all 128 channels. This revealed a significant time-frequency cluster (summed across significant sensors) in which alpha power (spanning frequencies in the beta range but with a peak at 9 Hz) was reduced for “Remember” vs. “Forgot” trials from ~800-1500 ms post stimulus onset (P_cluster_ < .001). The scalp topography of the effect (summed across significant time/frequency bins) indicated a widespread extent, with a slightly stronger effects at left compared to right hemisphere sensors.

**Figure S4.**
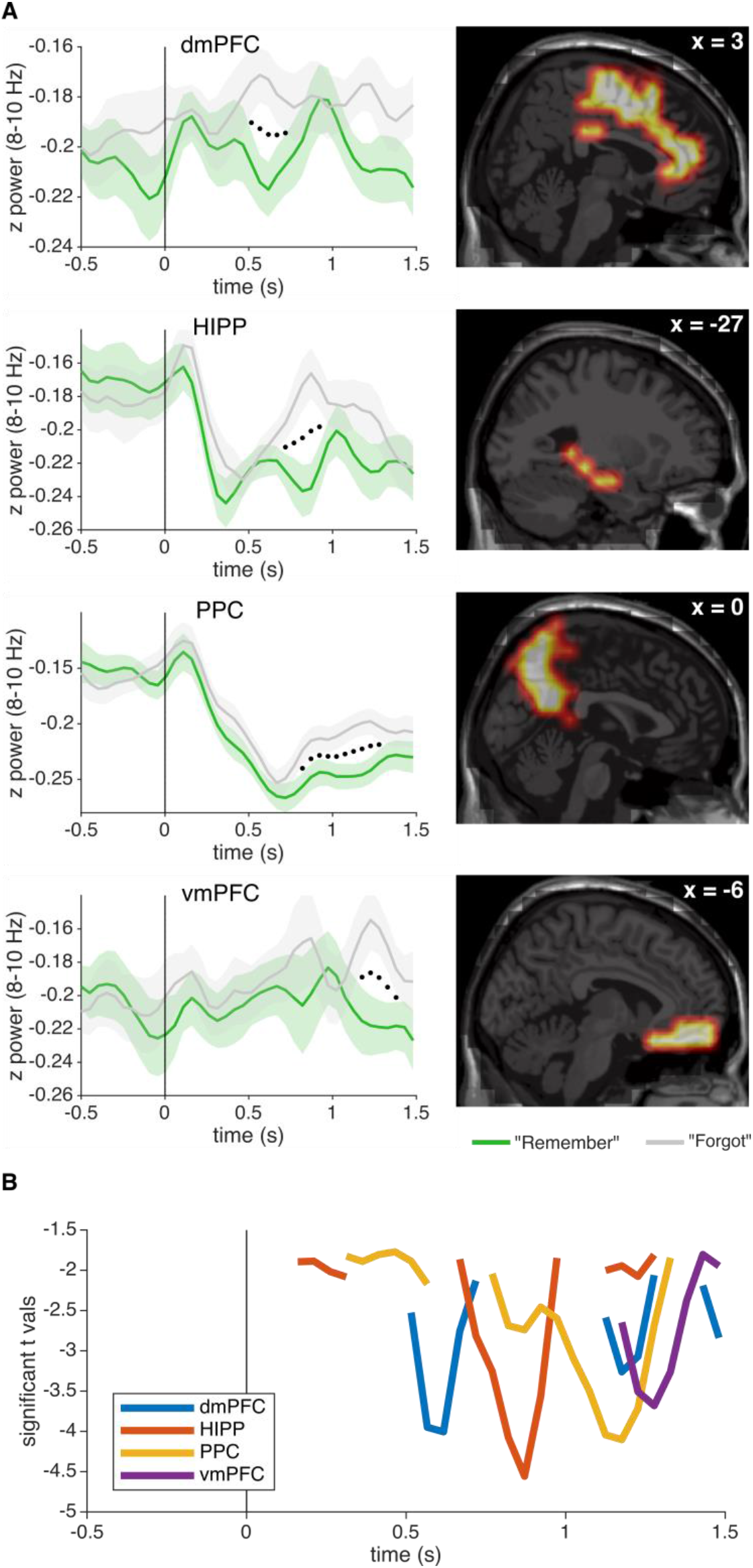
Temporal dynamics of EEG alpha (8-10 Hz) recall effects. **A.** *Left*: z power time courses for “Remember” (green) and “Forgot” trials (grey; mean +/− SEM of condition differences across participants). Black dotted line marks time points of significant condition differences surviving cluster-based correction for multiple comparisons across time. *Right*: Regions of interest defined as the overlap of significant voxels resulting from the main voxel × time analysis (cf. main Figure 4A) and AAL masks. X coordinates refer to MNI space. **B.** T values for the comparison of “Remember” vs. “Forgot” in each region, thresholded at P < .05 (uncorrected). dmPFC = dorsomedial prefrontal cortex/anterior cingulate cortex, HIPP = hippocampus, PPC = posterior parietal cortex, vmPFC = ventromedial prefrontal cortex.

